# First draft genome assembly and characterization of sponge *Halisarca dujardinii* reveals key components of basement membrane and broad repertoire of aggregation factors

**DOI:** 10.1101/2024.02.06.578935

**Authors:** Ilya Borisenko, Alexander Predeus, Andrey Lavrov, Alexander Ereskovsky

## Abstract

How features characteristic of multicellular animals emerged in evolution and how the body plan of particular taxa was shaped are hotspots of modern evolutionary biology. We can get closer to answering them by studying animals that occupy a basal position on the phylogenetic tree, such as sponges (Porifera). We sequenced the genome of the sponge *Halisarca dujardinii* using Oxford Nanopore and Illumina technologies and made an assembly of long reads, followed by polishing with short reads. The resulting assembly had a size of 176 Mb, matching the prediction from the k-mer distribution, and an N50 of about 785 Kb. By analyzing transposable elements in the genomes of *H. dujardinii* and a number of other sponges, we found that a significant portion of the genome (more than half for Demospongiae) is occupied by repeats, most of which are evolutionary young. RNA-seq data were used to predict about 14000 genes in the genome, several times less than in other Demospongiae. By analyzing ortholog groups unique to *H. dujardinii* among sponges and higher invertebrates, we found overrepresented genes related to the extracellular matrix. The extracellular matrix of *H. dujardinii* contains, among others, key basement membrane components such as laminin, nidogen, fibronectin, and collagen IV, for which phylogenetic analysis has confirmed that it belongs to this type of nonfibrillar collagen. In addition, we showed in *H. dujardinii* 14 aggregation factor genes responsible for cell recognition and adhesion. They are organized in a genomic cluster and have at least two types of domains: Calx-beta, responsible for calcium ion binding, and Wreath domain, unique for this type of molecules. Our obtained assembly and annotation will further expand the understanding of genome evolution at the emergence of animal multicellularity, and will serve as a tool to study the regulation of gene expression by modern methods.

## Introduction

The phylum Porifera occupies a basal position on the phylogenetic tree. Although recent studies challenge their sister relationship with the rest of the Metazoa, they have long been regarded as the first of the multicellular organisms (given the controversial position of Placozoa) ^1^. The phyla consist of four classes, each with characteristic features. The class Calcarea includes sponges with a well-developed skeleton composed of calcium carbonate. Sponges in the class Hexactinellida have a syncytial structure in adulthood. The class Demospongiae, or common sponges, is the largest, and includes sponges with a skeleton made of silicon oxide or organic substance (spongin) ^2^. The class Homoscleromorpha has recently been separated from the class Demospongiae, and includes sponges with the most advanced features of Eumetazoa: basement membrane contained collagen type IV and adherens-like intercellular junctions ^3–5^.

Over the last few years, a number of sponges have emerged that can be called model species, facilitated by remarkable features of their biology. For example, sponges have a huge number of obligate endosymbionts from among bacteria and archaea, therefore, much research has been devoted to the study of symbiosis ^6,7^. Sponges (and their endosymbionts) produce secondary metabolites, which often have high biological activity and are considered potential marine drugs ^8^. Finally, sponges have remarkable regenerative abilities: they can regenerate a small body fragment that has been removed, and the cellular mechanisms of this process differ radically between species^9^. In addition, when the sponge body dissociates into individual cells, they are able to come together and form a primmorph, a spherical conglomerate that develops into a new sponge ^9^.

The first sponge genome, of *Amphimedon queenslandica*, was published just 14 years ago ^10^. Then, with a large gap, new genomes began to appear, with assembly levels from shotgun to chromosome level (Belahbib et al., 2018 for *Oopsacas minuta* and *Oscarella lobularis*; Fortunato et al., 2012 for *Sycon ciliatum*; Francis et al., 2017 for *Tethya wilhelma*; Kenny et al., 2020 for *Ephydatia muelleri*; Santini et al., 2023 for *O. minuta*; Schultz et al., 2023 for two unidentified Cladorhizida and Hexactinellida sponges). And also, in the past year, eleven sponge genomes have been assembled to chromosome level based on PacBio HiFi and Hi-C data by the Aquatic Symbiosis Genomics Project (https://www.aquaticsymbiosisgenomics.org/).

Even within the class Demospongiae, the evolutionary distance between families is so great that their genome size and gene content can differ by several times (see below; Wörheide et al., 2012). Here we report an assembly of the 176-Mbp genome of demosponge *Halisarca dujardinii* as new sponge reference genome. During our work, we encountered difficulties in DNA purification, sequencing, and assembly. We optimized the isolation protocol and analyze the reasons for the mentioned difficulties. We report significant characteristics of the sponge genome, including the context of genes, transposable elements, analyzed gene families within the Porifera for orthology, and evaluated the repertoire of extracellular matrix proteins with a focus on basement membrane components and aggregation factors.

## Materials and methods

### Sample collections

Adult sponges *H. dujardinii* were collected in the area surrounding the Marine biological station «Belomorskaya» of Saint Petersburg University in the Kandalaksha Bay of the White Sea from May to September. Fucus algae with sponges were taken from the sublittoral horizon (1-5 m) using a small anchor and then placed in aquaria with circulating seawater and aeration at 12 degrees Celsius. Reproducing sponges were collected in June-July. Individuals with embryos were placed in cups with water and visually inspected several times a day for the emergence of larvae from the maternal organism. After larval emergence began, the cups were placed under a desktop lamp, and larvae were collected near the water surface. Larvae were collected manually by pipette and concentrated using a 40 µm mesh sieve. Body fragments of sponges and larvae were frozen in liquid nitrogen and stored there until nucleic acids were isolated. RNA was isolated from free-swimming larvae, late cleavage stages, and regenerates at stages 3, 12, and 24 h after injury. RNA was isolated using Lumisol (Lumiprobe, cat. no. 91415) and purified using spin columns (Evrogen, cat. no. BC033). DNA was isolated using the gravity flow Genomic Tip kit (Qiagen, cat. no. 10223) and by the phenol-chloroform method ^17^. The quality of nucleic acids was assessed by agarose gel electrophoresis (1% for RNA, 0.7% for DNA) and on-chip electrophoresis (Bioanalyzer 2100, Agilent).

### Sequencing

Oxford Nanopore libraries were prepared using the SQK-LSK109 (Oxford Nanopore Tech-nologies, ONT) protocol and sequenced on two MinION flowcells (FLO-MIN106) and one PromethION flowcell (FLO-PRO002) for 72 hours or until the working pore number fell below 50. The raw signal obtained as fast5 files was converted to fastq files, reflecting nucleotide sequence and signal quality, by the basecaller Guppy v5.0.11+2b6dbff in high-fidelity mode, using the con-figuration file “dna_r9.4.1_450bps_sup.cfg” (for MinION) and “dna_r9.4.1_450bps_sup_prom.cfg” (for PromethION).

For polishing, we used genomic DNA reads previously obtained by Maja Adamska and de-posited in the SRA under the number ERX1222365. DNA in this experiment was also isolated from larvae, libraries were prepared with TruSeq RNA V2, and reads were obtained on an Illumina HiSeq 2000 instrument.

The RNA library was prepared from the polyadenylated fraction, and sequenced on a NovaSeq6000 150 PE with yield of 25-35 million of reads per sample.

### Assembly

Illumina reads were processed with the BBduk software (v38.07, palindromic mode; options “ktrim=r k=23 mink=11 hdist=1 tpe tbo”) to remove adapter sequences. All Oxford Nanopore reads were also combined, and filtered using the program Filtlong v0.2.0 using the cleaned Illumina se-quences as a reference. We chose 5 popular assemblers for assembly: Flye v2.9-b1768, Canu v2.1.1, Shasta v0.7.0, Raven v1.5.1, and Wtdbg2 (Redbean) v2.5. For polishing we used the programs Ra-con, Polca and Pilon. To assess the quality of the assembly, we used the program busco 5.2.2 with the “--augustus” option to find *de novo* genes in the studied genomes. The “metazoan” collection (954 genes) from OrthoDB v10 was used as a database.

### Repeat analysis

Repetitive elements in the genome were analyzed using the Earl Grey pipeline ^18^, which includes RepeatModeler2, RepeatMasker v4.1.2, and LTR_Finder v1.07. This tool allows *de novo* identification of transposable elements (TEs), clustering them into families, classifying the families against the open Dfam database, and using the resulting library to mask repeats in the genome. Custom repeat libraries were created for the *H. dujardinii* genome, *Spongilla lacustris* (GenBank acces-sion GCA_949361645.1), *E. muelleri* (GenBank accession GCA_013339895.1), *O. lobularis* (GenBank accession GCA_947507565.1), *Aphrocallistes beatrix* (GenBank accession GCA_963281255.1), *O. minuta* (GenBank accession GCA_024704765.1) and *S. ciliatum* (https://doi.org/10.5061/dryad.tn0f3).

### Transcriptome assembly and gene prediction

Transcriptome data used for training and as evidence during gene prediction. Raw paired-end sequences of cDNA were quality filtered and trimmed using Trimmomatic v0.32 with default settings. Unpaired reads were discarded. Pooled reads of five samples (larvae, embryos, regenerates at 3, 12 and 24h) were assembled *de novo* using Trinity 2.13.2. Assembled transcriptome was cleaned by aligning reads of genomic DNA from larvae. The alignments were done using bowtie2 v2.4.4 with default parameters. Transcriptome contigs not aligned to any of the gDNA read were removed from the assembly. Contigs were clustered and merged with cd-hit-est v4.8.1 with identity threshold 0.95.

Another transcriptome was assembled in genome guided manner. The same five cDNA libraries were mapped to the genome using STAR v2.7.10a. Resulting .bam file was used as input for Trinity v2.15.1 in genome guided mode. Resulting contigs were also clustered with cd-hit-est with the same threshold.

Additional set of transcripts was generated with StringTie v2.2.1. Each cDNA library was independently mapped to the genome with STAR, and .bam file used as input for StringTie. Resulted .gtf were assembled with TACO v0.7.3 with default settings. These TACO-produced transcript coordinates provided to PASA pipeline together with *de novo* and genome guided transcriptomes ^19^, where only transcripts with more than 90% transcript coverage (parameter -- MIN_PERCENT_ALIGNED) and 95% identity (parameter --MIN_AVG_PER_ID) to the genome were merged.

Ab initio gene predictions were performed on the repeat-masked assembly with BRAKER v3.0.6 pipeline based on two different programs — Augustus v3.5.0 ^20^ and Genemark-ES v2.3e ^21^. The same five cDNA libraries mapped to the genome were used for training and as evidence in gene prediction.

Gene evidence and predicted gene structure were combined using EVM ^19^. The evidence included a) ab initio predictions generated by Augustus and GeneMark, b) PASA generated consensus transcript assemblies based on *de novo* assembly, genome guided assembly and StringTie transcripts, and c) the miniprot v0.12 generated assembly of UniProtKB/Swiss-Prot (2023-11-08) to the genome. The evidence weights for EVM were set as follows: transcripts assembled by PASA – 10, miniprot alignment – 2, *ab initio* genes prediction – 1. UTRs were added onto the gene models predicted by EVM by two sequential PASA rounds including annotation loading, annotation comparison and annotation updates to maximize incorporation onto gene models predicted by EVM, as per the suggestion of the authors in the PASA pipeline manual (http://pasapipeline.github.io/). Gene models together with assembly, predicted proteins and Trinotate report are deposed in Github repos-itory https://github.com/mezoderma/Hdu-genome.

### Gene annotation and orthology analysis

Open reading frames (ORFs) for all genes were predicted using Transdecoder v5.7.1. All best ORF candidates were analysed for protein domains, signal sequences and transmembrane domain using hmmer 3.0, signalp 6, diamond v2.0.15.153 and tmhmm 2.0, and combined to annotate each ORF using Trinotate v4.0.2. Protein sequences for the ortholog search were downloaded from the corresponding genomic projects in GenBank indicated in section “Repeat analysis” (for *E. muellery*, *O. lobularis*, *O. minuta*, *Trichoplax adhaerans, Nematostella vectensis*, *Branchiostoma floridae*) and from https://doi.org/10.5061/dryad.tn0f3 (for *S. ciliatum*, Fortunato et al., 2012). Orthogroups were constructed using OrthoFinder v2.5.5, and analysis results were visualized using the UpSetR library for R. GO enrichment performed with GOseq v1.54.0; GO terms above threshold 0.0001 were visualized with REVIGO ^22^.

## Results and discussion

### DNA Extraction and Sequencing

The adult sponge *Halisarca dujardinii* (Fig. 1A) contains a huge number of cells per unit volume of tissue, so it is a fruitful target for genomic DNA isolation. Its mesohyl is rich in glycopro-tein-rich extracellular matrix. At first glance, the large concentration of polysaccharides did not prevent the isolation of high-molecular weight genomic DNA of sufficiently high quality. Thus, we ob-tained DNA with an average molecular weight of more than 150 kb and A260/A230 > 2.1 from both cells and whole sponges with classical phenol-chloroform purification (Sambrook et al., 2001 after Blin and Stafford, 1976). In the former case, the whole sponge was homogenized and subjected to lysis, while in the latter case, it was preliminary dissociated in artificial calcium-free seawater. Intercellular contacts in sponges are calcium-dependent, so the removal of calcium with EDTA leads to the separation of their tissues into individual cells that remain viable. After dissociation, the cells were collected by centrifugation, washed of the extracellular matrix, and then lysed.

**Fig. 1.**
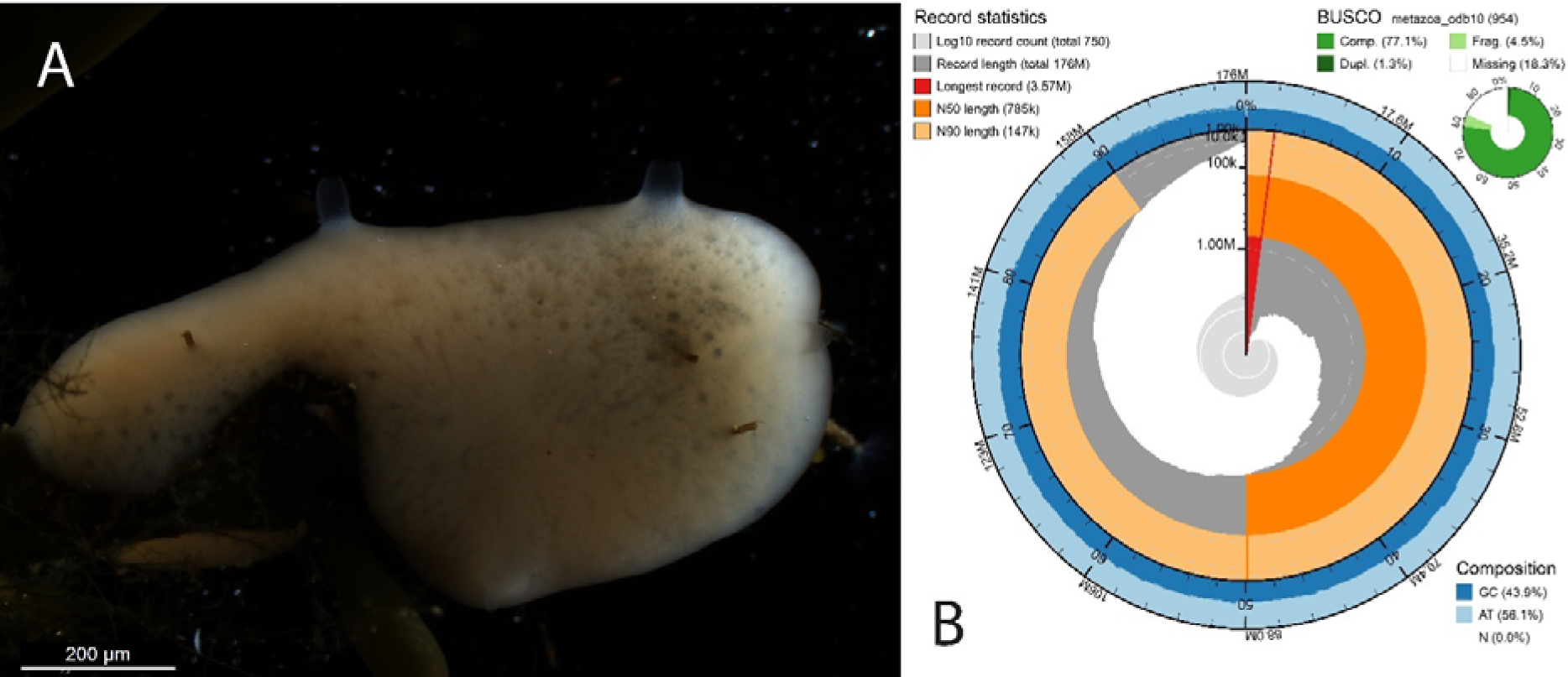
Demosponge *H. dujardinii*. (A) Adult animal. (B) The snail plot shows that the final version of the *H. dujardinii* assembly has N50 of 785 Kb, the longest scaffold is 3.57 Mb long.

In pilot sequencing experiments on MinION, we used the Rapid Sequencing kit (SQK-RAD004), which allows to obtain the maximum read length. However, in two cell experiments involving a positive control, we found that the sequencing of genomic DNA was apparently interfered with by some substance accompanying the purified DNA. Also, adapter ligation did not occur during library preparation for Illumina, from which we hypothesize that an unknown substance interferes with ligase activity during library preparation for both ONT and Illumina.

In sponges, a large number of secondary metabolites with different biological activities are known ^24^. At least some of these probably serve a protective function for the immobile sponge, whose defense capabilities are severely limited. Assuming that the interfering sequencing agent may be a secondary metabolite of the adult sponge or associated with the extracellular matrix, we isolated genomic DNA from larvae. The swimming larva of *H. dujardinii* does not contain an extracellular matrix and probably does not produce secondary metabolites prior to metamorphosis.

Genomic DNA from larvae was successfully sequenced on ONT and Illumina in several rounds. In the first round, we compared purification using phenol-chloroform and chromatography on gravity-flow columns (QIAGEN Genomic-tip kit), and chose chromatography, which allowed us to avoid toxic phenol, obtaining DNA lengths up to 100 kb. In ONT, we have favored 1D chemistry for library preparation, which, compared to Rapid ligation kit, allows us to obtain shorter read lengths but a larger amount of data. Sequencing on two MinION cells and one PromethION cell resulted in 26.3 GB of reads. Using only reads that passed the pass/fail filter, we kept 90% of the longest reads, yielding 19 GB of reads with an N50 of about 17 kb after the filter. The data was de-posed in GenBank under BioProject PRJNA1055929.

Contamination of genomic DNA by reads originating from inevitably present microorganisms and viruses is often a significant source of problems in the assembly, annotation, and analysis of eukaryotic genomes. We analyzed the taxonomic composition of Illumina reads using Kraken v2.1.2 software and the databases nt (version dated 01/27/2020) and k2_pluspf_20210517 (version dated 05/17/2021; including *Homo sapiens* and all bacteria, archaea, viruses, and yeasts known to RefSeq). Analysis of Oxford Nanopore reads using Kraken2 is unreliable due to the high error rate in the reads. The result showed little contamination with other species: only 0.01% of the reads were attributed to archaea, 1.10% to bacteria, 0.20% to yeasts, 0.50% to unicellular algae, and 0.13% to viruses. Thus, contamination of the study samples with microorganisms is minimal and will not interfere with genome assembly and annotation.

### Genome Size Estimation

Illumina reads were used to estimate genome size (ONT reads contain too many errors for such an analysis). There are several ways to estimate genome size, both experimental (based on DNA content in the nucleus) and based on genomic reads (pre- and post-assembly). Estimation of genome size and properties before assembly based on k-mer distribution is, *de facto*, a common step in genome assembly.

We used the actively supported software GenomeScope 2.0, which allows us to obtain estimates of genome size, degree of heterozygosity, and number of repeats present. The good (approximately 200-fold) coverage of the genome by Illumina reads allowed us to obtain a stable model that does not change when the size of the k-mer used changes from 17 to 27. The estimation results for k=21 is given in Supp. Fig. 1 and Supp. Table 1. We determined that the haploid genome size is 188 Mb.

**Table 1.** The results of assembling Oxford Nanopore reads by different assemblers.

| Assembler | Options | N50, bp | Genome length, bp | BUSCO (954 metazoan genes, odb10) |  |  |  |
| --- | --- | --- | --- | --- | --- | --- | --- |
|  |  |  |  | Complete | Duplicated | Fragmented | Missed |
| Flye | - | 176,796 | 342,205,990 | 737 | 64 | 38 | 179 |
|  | min_overlap=8000 | 182,801 | 352,750,373 | 734 | 85 | 37 | 183 |
|  | min_overlap=10000 | 176,642 | 358,772,653 | 731 | 80 | 38 | 185 |
| Canu | - | 156,289 | 502,417,763 | 735 | 281 | 32 | 187 |
| Shasta | - | 746,535 | 176,219,426 | 706 | 13 | 57 | 192 |
|  | minReadLength 2000 | 798,451 | 178,626,056 | 713 | 13 | 49 | 192 |
|  | minReadLength 5000 | 976,698 | 178,393,878 | 720 | 11 | 51 | 183 |
|  | minReadLength 8000 | 917,022 | 180,849,520 | 708 | 14 | 56 | 190 |
| Raven | - | 234,518 | 245,632,875 | 715 | 37 | 51 | 188 |
| Wtdbg2 | - | 450,332 | 294,732,264 | 703 | 25 | 46 | 205 |

A recent publication describes a preliminary and unpublished assembly of the *H. dujardinii* genome, estimating a genome size of 194 Mb based on 44-fold coverage of Illumina reads ^25^. This estimate was made using the Kmergenie program. The recently assembled genome of a closely re-lated species, *Halisarca caerulea*, published in GenBank under accession GCA_963170055.1, is estimated to be 195.7 Mb. Results obtained in different laboratories by different methods showed a remarkable similarity in the estimation of haploid genome size. Heterozygosity of the genome was estimated at 0.62%; it should be noted, however, that this estimate is based on k-mers and is not capable of estimating heterozygosity of repeat regions.

### Genome Assembly

Given the recommendations for hybrid assembly, the high coverage (about 100x Oxford Nanopore and about 200x Illumina reads) conditioned the chosen strategy, namely assembling the genome from long reads, followed by polishing the genome with Illumina reads. We chose 5 popular assemblers for assembly: Flye v2.9-b1768, Canu v2.1.1, Shasta v0.7.0, Raven v1.5.1, and Wtdbg2(Redbean) v2.5. The results of the assemblies are presented in Table 1.

One can easily see from the above values that the assembly is problematic: the total length of the haploid assembly differs between some programs by three times. Evaluation with the busco software shows that a significant part of the classical single-copy orthologs characteristic of animals is missing from the assembly. We chose the assembly made by the program Shasta with a minimal read length of 5000 bp, which has the highest Benchmarking Universal Single-Copy Orthologue (BUSCO) level.

We can only assume one scenario given the high taxonomic purity of the reads obtained, the high coverage of both long and short reads, and the low estimate of heterozygosity obtained by k-mer analysis: the *H. dujardinii* genome contains a significant number of evolutionarily young re-peats that are present in a highly heterozygous state. This is indirectly evidenced by the diagram of contig coverage distribution obtained by Flye assembly (Supp. Figure 2): we can see a bimodal distribution, as well as a significant number of poorly covered contigs, which are assembly or alignment artifacts.

We applied long-read polishing using the Racon v1.4.20 software to improve the resulting assemblies. Nine rounds of iterative polishing were performed. Unfortunately, there was little improvement in the assemblies; after a slight improvement in the first round of polishing, BUSCO values fluctuated around some equilibrium value (Supp. Figure 3).

Polishing with Illumina short reads improved the assembly to some extent. We tried sequential polishing with Racon, Polca, and Pilon software. An increase in BUSCO values was observed after the first round but decreased in subsequent iterations. The BUSCO value for the *H. dujardinii* assembly was 77% for complete gene models, and with the inclusion of partial sequences, the genome accounted for 81.7% of the metazoan reference gene set. The size of the assembly after polishing was 176 Mb, N50 value was about 785 Kb, and it contains 746 contigs (Fig. 1B).

### Repetitive elements analysis

Studies focusing on genome evolution are based mainly on the comparison of protein-coding genes: on the search for families common to different taxa, lineage-specific expansion or loss. At the same time, it is known that in the majority of Metazoa, the transposable elements (TEs) occupy a significant part of the genome and are actively involved in genome evolution. Only a few studies have analyzed TEs of the sponge genome ^14,15,26^. The diversity and role of these genetic elements in such ancient multicellular organisms as sponges deserve more attention.

The TE repertoire of four sponges*, E. muelleri, T. wilhelma, A. queenslandica*, and *S. ciliatum*, was analyzed using *de novo* repeat modeling ^14^. Repeats have been shown to occupy a substantial portion of the sponge genome (29 to 49%), and most of them are currently unclassified. It is noticeable that the expansion of different classes of repeats occurs specifically within different taxa. Kenny et al. provide a logical hypothesis that genome size correlates with the quantity of re-peats in the genome, and the proportion of repeats is higher in larger genomes ^14^. Interspersed re-peats represent 34% of the *O. minuta* genome, and these are mostly represented by unclassified re-peats (18%) and DNA transposons (15%; Santini et al., 2023). The *A. queenslandica* genome contains 43.2% of repetitive elements, most of which are DNA transposons (4.16%). Among these non-autonomous DNA transposons, two miniaturized transposable elements, MITEs, named Queen1 and Queen2, have been described ^26^. They probably originate from the Tc1/Mariner-like repeats family. Usually MITEs are low-copy numbers, but in *A. queenslandica*, they are abundant in the genome (3800 and 1700 repeats, respectively), and Queen1/Queen2 have not been detected in the genomes of other animals. MITEs can introduce new splice sites, providing domain shuffling and inducing the formation of new multidomain proteins.

We analyzed the diversity of TEs in the genomes of a number of sponges belonging to all four classes: Demospongiae (*H. dujardinii*, *E. muelleri*, and *S. lacustris*), Calcarea (*S. ciliatum*), Homoscleromorpha (*O. lobularis*), and Hexactinellida (*A. beatrix* and *O. minuta*). TEs were identified *de novo* and classified by comparison with the Dfam database using a new pipeline based on RepeatModeler, Earl Grey (Supp. Table 2). Overall, repeats span from 23 to 59% of the genome in sponges (Fig. 2). In the genome of *H. dujardinii*, TEs occupy 53.67% of the genome. The *de novo* library included 7450 families of repeats. Most of them (approximately 2/3, or 35.7% of the genome) are represented by repeats that could not be assigned to any described family. The identified TEs of *H. dujardinii* are dominated by retrotransposons of the SLACS CRE and BovB families belonging to LINE, and DNA transposons of the Tourist and Harbinger superfamilies, as well as the Pogo superfamily of the IS630-Tc1-mariner group.

**Fig. 2.**
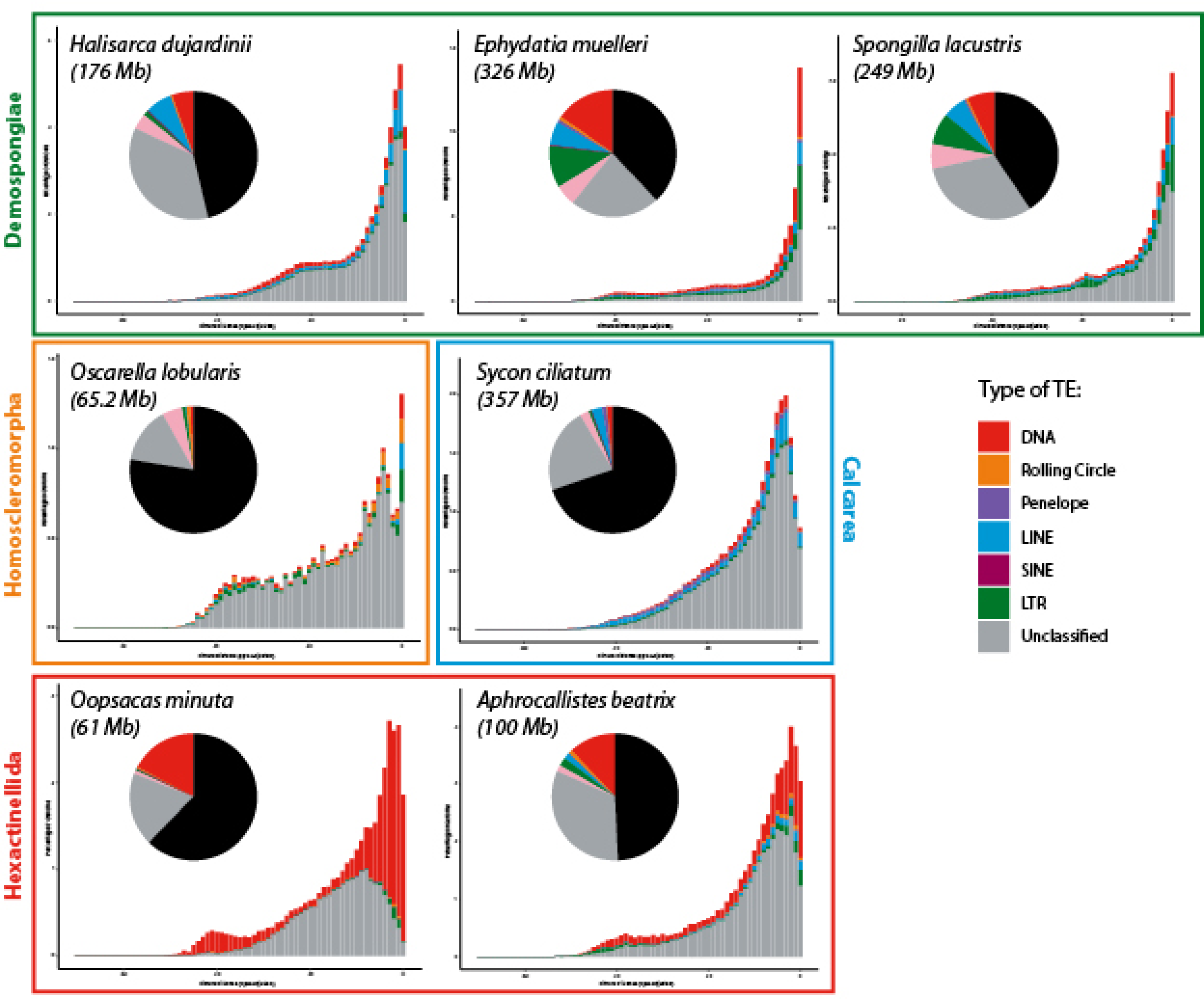
Repeat landscape plots illustrating TE accumulation history for genome of sponges from different classes, based on Kimura distance-based copy divergence analyses, with sequence divergence (CpG adjusted Kimura substitution level) illustrated on the x-axis, percentage of the genome represented by each TE type on the y-axis, and transposon type indicated by the color chart on the right-hand side. Major TEs: LINE, long interspersed nuclear element; LTR, long terminal repeat; Penelope, family of retrotransposons; Rolling Circle, family of Helitron transposons; SINE, short interspersed nuclear element.

**Table 2.**
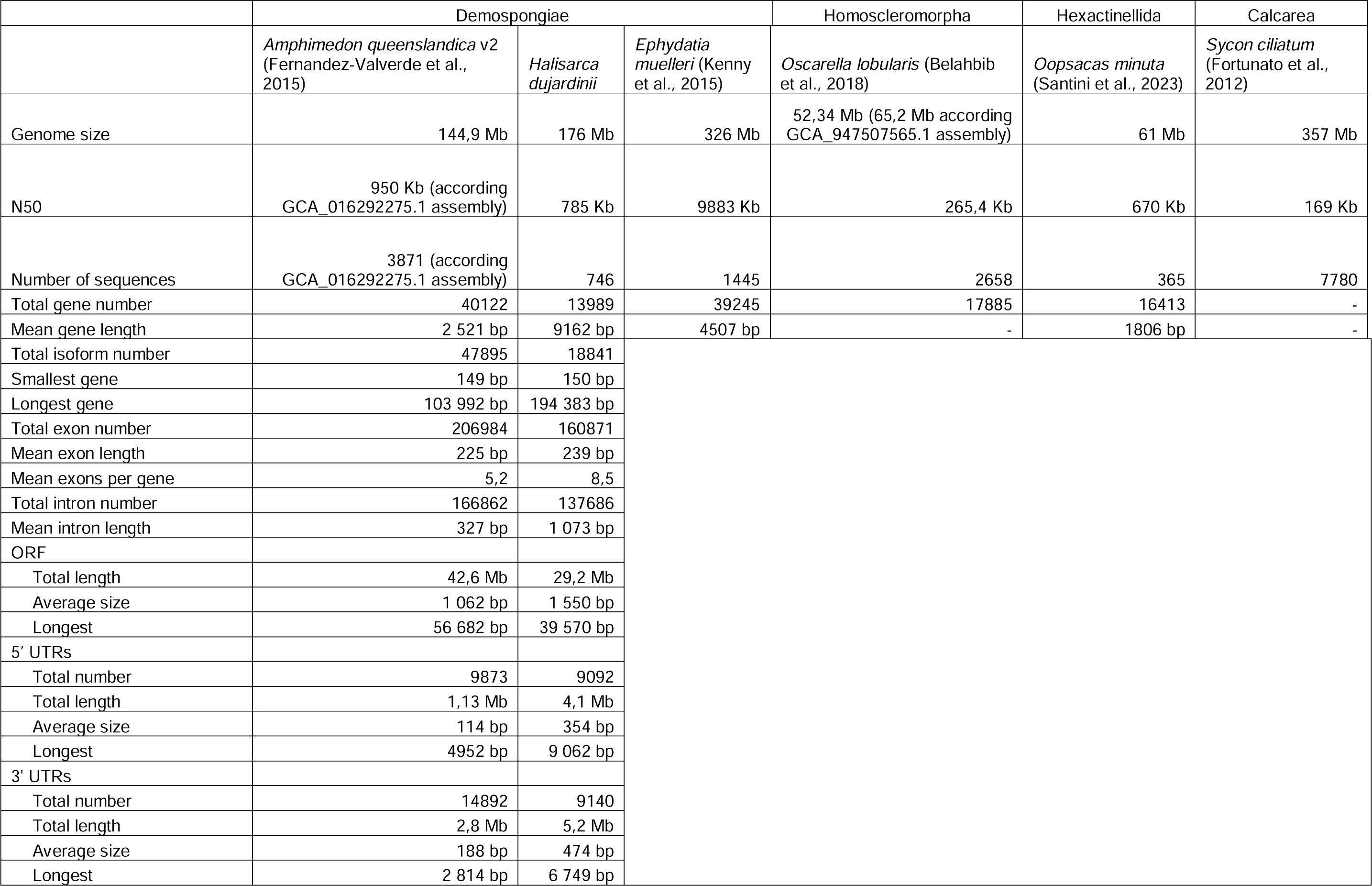
Genome assembly statistics for different sponge species.

**Table 3.** Comparison of completeness and composition of two *H. dujardinii* predicted gene models.

|  | *H. dujardinii* transcriptome assembly, v1<br>(GenBank HADA00000000.1) | Genome-derived gene models |
| --- | --- | --- |
| # of genes | 105862 | 13989 |
| # of transcripts | 138992 | 18841 |
| # of ORFs | 40754 | 18841 |
| BUSCO | C:90.9%[S:16.9%,D:74.0%],F:2.5%,M:6.6%,<br>n:954 | C:85.8%[S:84.1%,D:1.7%],F:4.3%,M:9.9<br>%,n:954 |
| Metazoa BBH | 14273 | 9803 |
| Non-metazoa BBH | 1648 | 909 |
| No blast hit with PFAM | 7827 | 3074 |
| No blast hit or PFAM | 17006 | 5055 |

In general, we were unable to describe any taxon-specific expansion of a particular class of repeats in individual sponge species or classes. It can be seen that different species show dominance of different groups and even superfamilies of repeats. For example, class Hexactinellida shows the prevalence of DNA transposons, whereas demosponges and calcareous sponges are dominated by retroelements like LINE and LTR. The genome of *O. lobularis* is generally very poor in repeats (22.28%). The genome of *O. minuta* is almost devoid of retroelements. SINEs were not identified in a number of sponge species (*Aphrocallistes*, *Oscarella*, *Ephydatia*, and *Spongilla*). At the same time, it should be taken into account that a substantial part of TEs remains unclassified, from 14 to 35%, and further studies are needed to elucidate the complete pattern of TE distribution within Porifera.

The interspersed repeat landscape demonstrates very recent (possibly ongoing) expansion of TEs in sponges (Fig. 2). The plot shows how many copies of each repeat are present in the genome as a function of the Kimura distance, which represents the genomic distance between the repeat and the consensus sequence. The distance increases with repeat divergence, *i.e.*, more recent repeats have a small Kimura distance. Peaks show large groups of copies of repeats with the same divergence and indicate a major expansion event of these elements. Broad peaks with a high Kimura dis-tance are characteristic of ancient TEs, and sharp peaks with a small distance are characteristic of young repeats. All sponges studied in this work are characterized by a sharp peak at a small Kimura distance, with substitution level ranging from 1 to 4. Most species have a second small peak between 20 and 42; this peak is absent in *S. ciliatum*.

### Evidence-based protein-coding gene annotation

To obtain the most complete set of expressed genes, we sequenced polyadenylated RNA from larvae, regenerates, and embryos at different developmental stages. We used RNA from an animal containing embryos or undergoing regeneration to assemble the transcriptome, as this can activate genes that are silent in the intact adult animal. A total of 5 libraries amounting to 63,9 Gb, corresponding to 330-fold ORF coverage, were used to assemble the transcriptome. The reads have been deposited in GenBank under the accession numbers SRR27694415, SRR27694424, SRR27709531, SRR27709534 and SRR27709540.

For annotation, we modified the strategy described for *Amphimedon* (Fernandez-Valverde et al., 2015; Fig. 3A). Thus, we combined using PASA the representative *de novo* assembly, genome-guided Trinity assembly, and independent genome-guided assembly for different stages generated with StringTie. The addition of StringTie-generated transcripts strongly improved the assembly: when it was included in the PASA pipeline, BUSCO was elevated from 70% to 88.8% (complete plus fragmented). To predict gene models, we used BRAKER3, an Augustus and GeneMark-ES-based pipeline that automatically performs model training and uses RNA/protein data as evidence. We compared models generated using RNA-seq and protein data (the longest ORFs isolated from the *de novo* transcriptome) and found that using RNA-seq data yielded a more complete set of BUSCOs (90.1% for RNA-seq, 86.5% for proteins; complete plus fragmented BUSCOs). A nearly identical BUSCO value (90.1%) was determined for the *E. muelleri* chromosome size assembly made with PacBio and Hi-C ^14^.

**Fig. 3.**
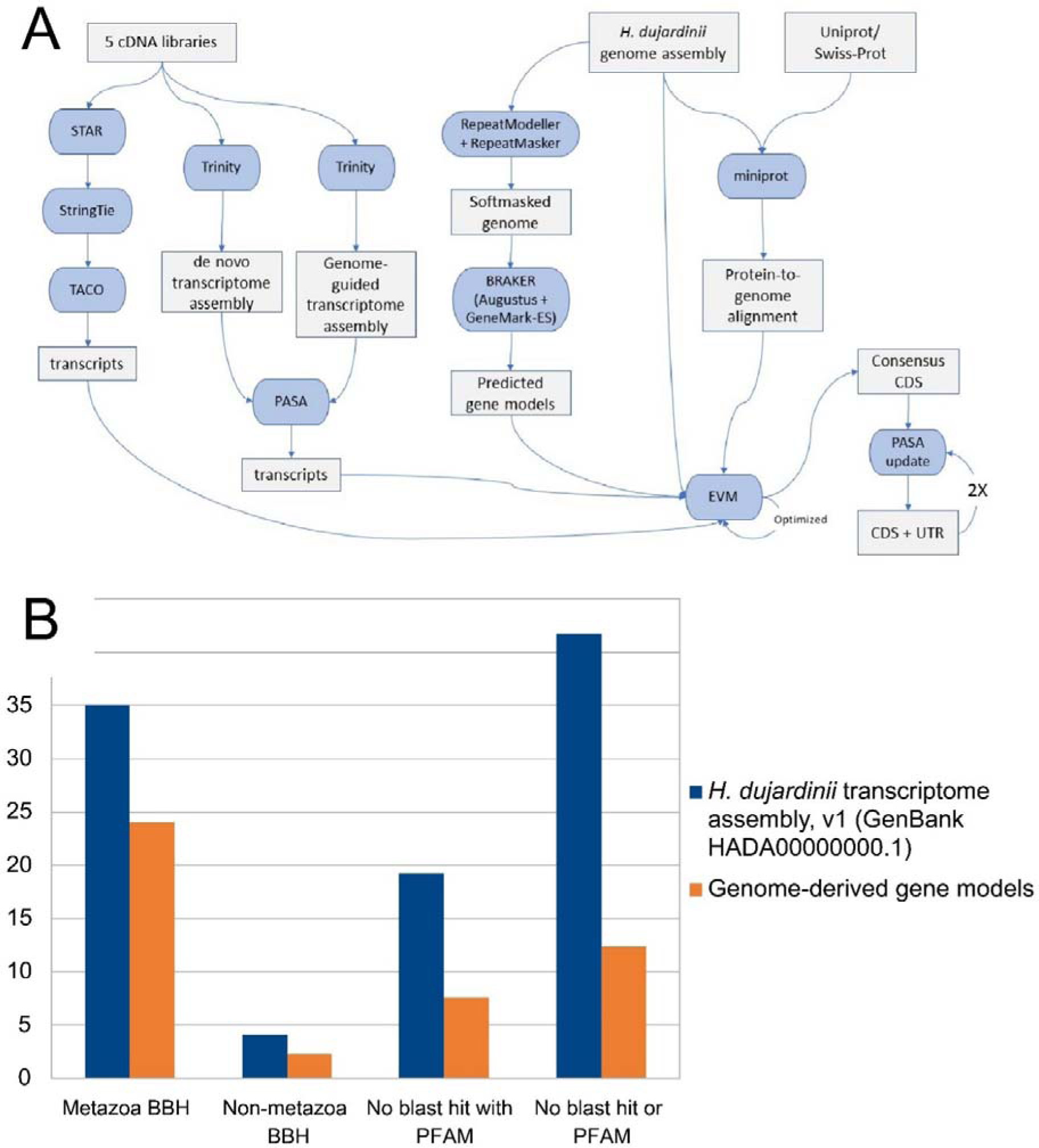
(A) Annotation pipeline. Combined data of RNA-seq generated from embryo, larva and regenerates of *H. dujardinii* were used to annotate the genome. This resulted in the annotation of CDS and UTRs. (B) BLASTP BBH annotation comparison. The y-axis shows the percentage of the total number of predicted transcripts. Also shown are proteins that have no significant blast hit yet contain an identifiable Pfam domain, and proteins with no significant blast hit and no identifiable Pfam domain.

All assembled transcripts, gene models, and protein-to-genome alignment were combined using EVM to predict protein-coding gene models ^19^.

The completed set of *H. dujardinii* genes contains a total of 13989 transcripts, which includes alternatively spliced gene isoforms that resulted in 18841 mRNAs. This is a relatively small number of genes for known sponge genomes (Table 2). In our annotation, 44% (8251) of all RNAs have both UTRs, and 53% (9981) of RNAs have at least one UTR. This is higher than in the last published annotation of *Amphimedon* genome, which was the best among sponges (37% of RNAs have at least one UTR; Fernandez-Valverde et al., 2015).

### Orthology analisys

We compared the predicted proteins of *H. dujardinii* with the proteomes of sponges and some other Metazoa using OthoFinder, a tool for finding orthologs and grouping them into groups (orthogroups). In addition to *H. dujardinii*, we took *A. queenslandica*, *E. muelleri*, *O. lobularis*, *S. ciliatum*, and *O. minuta* for analysis. First of all, we can see that the number of orthogroups in a species correlates with genome size (Fig. 4A). Thus, the highest number of orthogroups is characteristic of *E. muelleri* and *S. ciliatum* with relatively large genomes (about 10 and 12 thousand orthogroups, 326 and 357 Mb, respectively). The smallest number of orthogroups is in *O. minuta* with a compact genome (about 7 thousand orthogroups, 61 Mb). It is also noticeable that the largest number of orthogroups are unique for each species or common for all analyzed species. Thus, the largest number of orthogroups in the analyzed dataset (4554) belongs to *S. ciliatum*; they do not contain sequences of any other species. *O. minuta* stands apart, with species-specific proteins united into only 622 orthogroups, while the orthogroups common to other analyzed sponges taxa are slightly larger, 722 orthogroups. Taken together, this suggests that with the genome expansion, new (or divergence of existing) species-specific genes rather than duplication and increase in the number of orthologs in orthogroups occur.

**Fig. 4.**
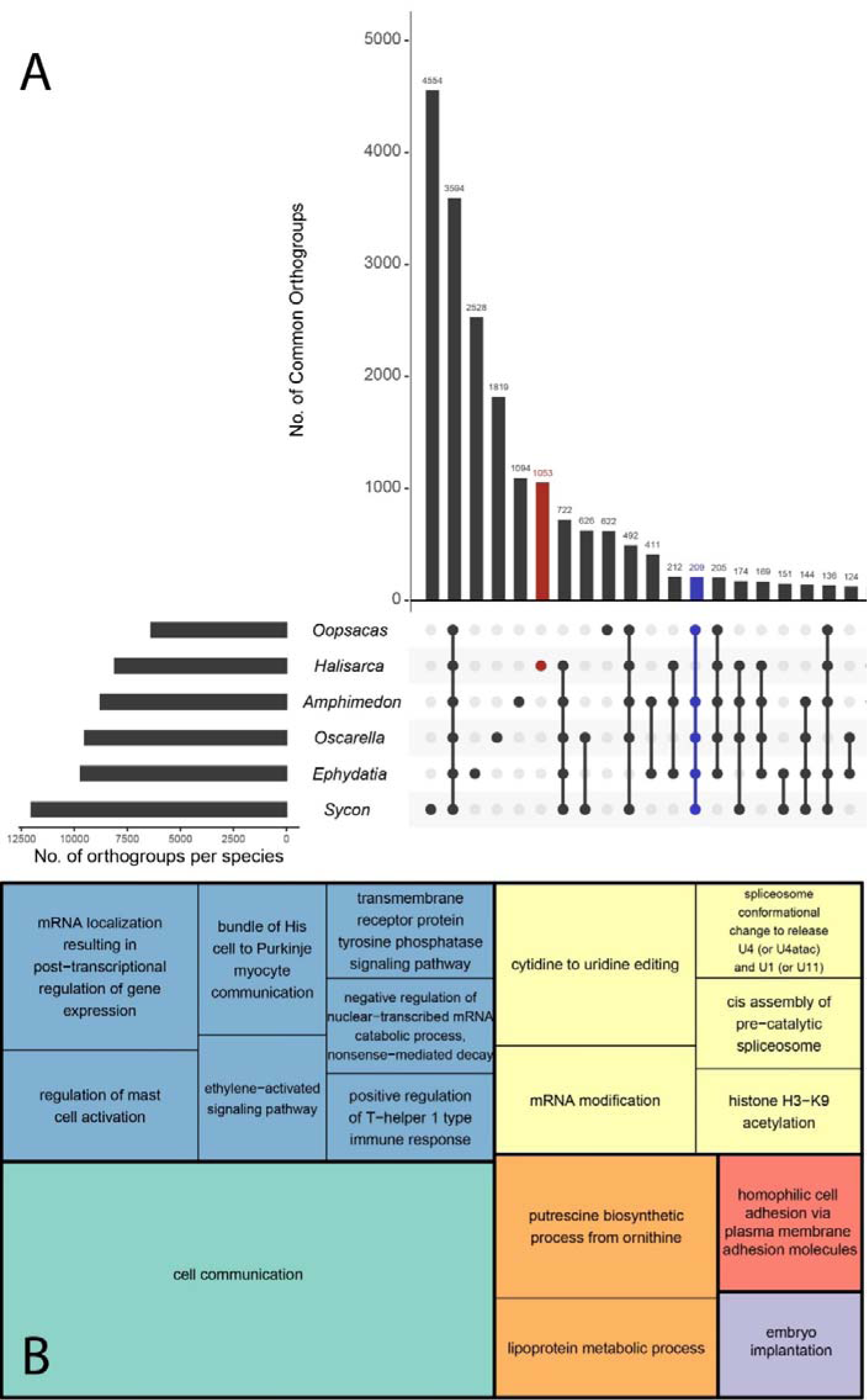
(A) Orthology analysis performed on six species representing all 4 classes of sponges. Each row under the bar chart corresponds to a species. The left side shows how many orthogroups the proteins of a given species represent. On the right is a matrix showing, using dots and lines connecting them, the sequences of which species are included in the orthogroups. The bar chart shows how many orthogroups have been found involving this set of species. Orthogroups of sequences unique to *H. dujardinii* are highlighted in red. Orthogroups that include all sponges except *H. dujardinii* are shown in blue. (B) GO enrichment of sequences unique for *H. dijardinii*. GO terms in category “Biological process” are indicated inside cells, and cell square proportional to p-value.

We then added representatives of other phyla, *Trichoplax adhaerens* (Placozoa), *Nematostella vectensis* (Cnidaria), and *Branchiostoma floridae* (Chordata) to the analysis to assess Porifera-specific gene expansion (Suppl. Fig. 4A). Again, orthogroups unique to the species or common to all species analyzed were the most represented. The number of species-specific orthogroups is more than 2-fold reduced in *Trichopax* compared to *O. minuta*, which has the minimum among sponges (251 vs 581). The addition of taxa did not change the number of orthogroups unique to *H. dujardinii*.

The 4511 sequences unique to *H. dujardinii* are clustered into 1053 orthogroups. 22% of these proteins had a BBH in Uniprot, and 44% had at least one domain annotated in Pfam. 54% of the proteins in these lineage-specific orthogroups did not have any annotation. We performed GO enrichment analysis of these proteins and found enrichment for the terms “Cell communication” (GO:0007154 in the Biological process category) and “Collagen-containing extracellular matrix” (GO:0062023 in the Cell compartment category) (Fig. 4B, Suppl. Table 3).

The group of proteins associated with the term “Cell communication” contained 80 peptides. Interestingly, 60 sequences transcribed from 25 genes were located at contig 20. Of these 80 proteins, 30 had BBHs with Uniprot, and all of these BBHs were with Metazoa sequences, so we can exclude a contamination artifact. Most of these sequences from contig 20 belong to aggregation factor family (see below).

### Extracellular matrix genes analysis

Manual inspection of sequences in both detected groups (“Cell communication” and “Collagen-containing extracellular matrix”) showed the presence of proteins somehow related to the extracellular matrix (ECM). Most of the proteins in the “Cell communication” group had the Calx-beta domain, a beta-folding domain consisting of tandem repeats and binding calcium ions. It has been described in deuterostomian transmembrane Na/Ca exchangers and in a number of extracellular matrix proteins (*e.g.*, extracellular matrix organizing protein FRAS1, FREM1/2). FRAS1 and FRAS1-related proteins show BBH for many extracellular matrix proteins due to the Calx-beta domain. An extracellular matrix protein, ECM3, was originally described to enable the migration of mesenchymal cells in blastocoel in the sea urchin ^29^. It had five Calx-beta domains, NG2-like (chondroitin sulfate proteoglycan), and transmembrane domains. The authors noted that the primary structure of ECM3 was similar to the sequence of MAFp4, an aggregation factor from demosponge *Microciona (Clathria) prolifera* ^30^. Later, a search for mutations responsible for the development of Fraser syndrome, a multisystem malformation usually comprising cryptophthalmos, syndactyly, and renal defects, resulted in the characterization of the FRAS1 protein gene in mammals. The FRAS1 protein sequence showed similarity to sea urchin ECM3, but in addition contained furin-like domains and von Willebrand factor type C domains ^31^. Due to these similarities, the gene for the ECM3 protein was named FREM – FRAS1-related extracellular matrix protein ^32^. The relationship between FRAS1 of mammals and FREM1/2 of echinoderms remains to be resolved, but it is clear that the group of sponge aggregation factors is monophyletic and not related by origin ^27^. The similarity between the sequences of FRAS-related proteins and AF can be traced due to the long length of conserved Calx-beta domains.

To better understand the composition of the extracellular matrix, we extracted the lines containing the term “extracellular matrix” from the Trinotate report and manually examined the resulting list for BBH and domain composition. Thus, we found orthologs with a characteristic set of domains of such structural proteins of the extracellular matrix as Porifera-specific fibrillar collagen ^33^ and fibrillar collagens of different classes, fibrinogen domain-containing proteins, cartilage matrix protein, mucin, as well as some proteins characterized but not having their own names (Sushi, nidogen, and EGF-like domain-containing protein, or SNED1). More than 100 intermolecular interaction partners have been predicted for this SNED1 protein, including integrin receptors and fibronectin, with which it partially immunolocalizes ^34^. SNED1 is widely expressed in mammalian embryos, especially intense in cells undergoing epithelio-mesenchymal transition ^35^. Cartilage matrix protein consists of two vWFA and EGF-like domains and has the ability to bind to collagen ^36,37^. A membrane-associated protein with a high level of glycosylation, Mucin, forms the backbone of mucus, covering the apical surface of epithelial cells in higher animals. Its functions are not limited to barrier function: mucins are able to regulate cell behavior and interactions ^38,39^. The enzymes procollagen lysine hydroxylase, glycosyltransferase and peroxidasin have also been found to provide post-translational modifications of collagens (Supp. Table 4). Many proteins with immunoglobuline or EGF domains are detected, but without reliable homology with any sequences from Uniprot.

We also specifically searched for genes of basement membrane components. Type IV collagen, laminin, nidogen (entactin), fibronectin, and the proteoglycan heparan sulfate ^40^ are characteristic for the basement membrane of Metazoa epithelia. Cells (including epithelial cells) express different types of integrins, which are transmembrane receptors that bind a wide range of ligands: fibronectins, collagens, laminins, fibrinogens, and others. We found gene orthologs of all basement membrane component proteins and integrin receptors (Supp. Table 4). All of them contain domains characteristic of Bilateria orthologs (Fig. 5A).

**Fig. 5.**
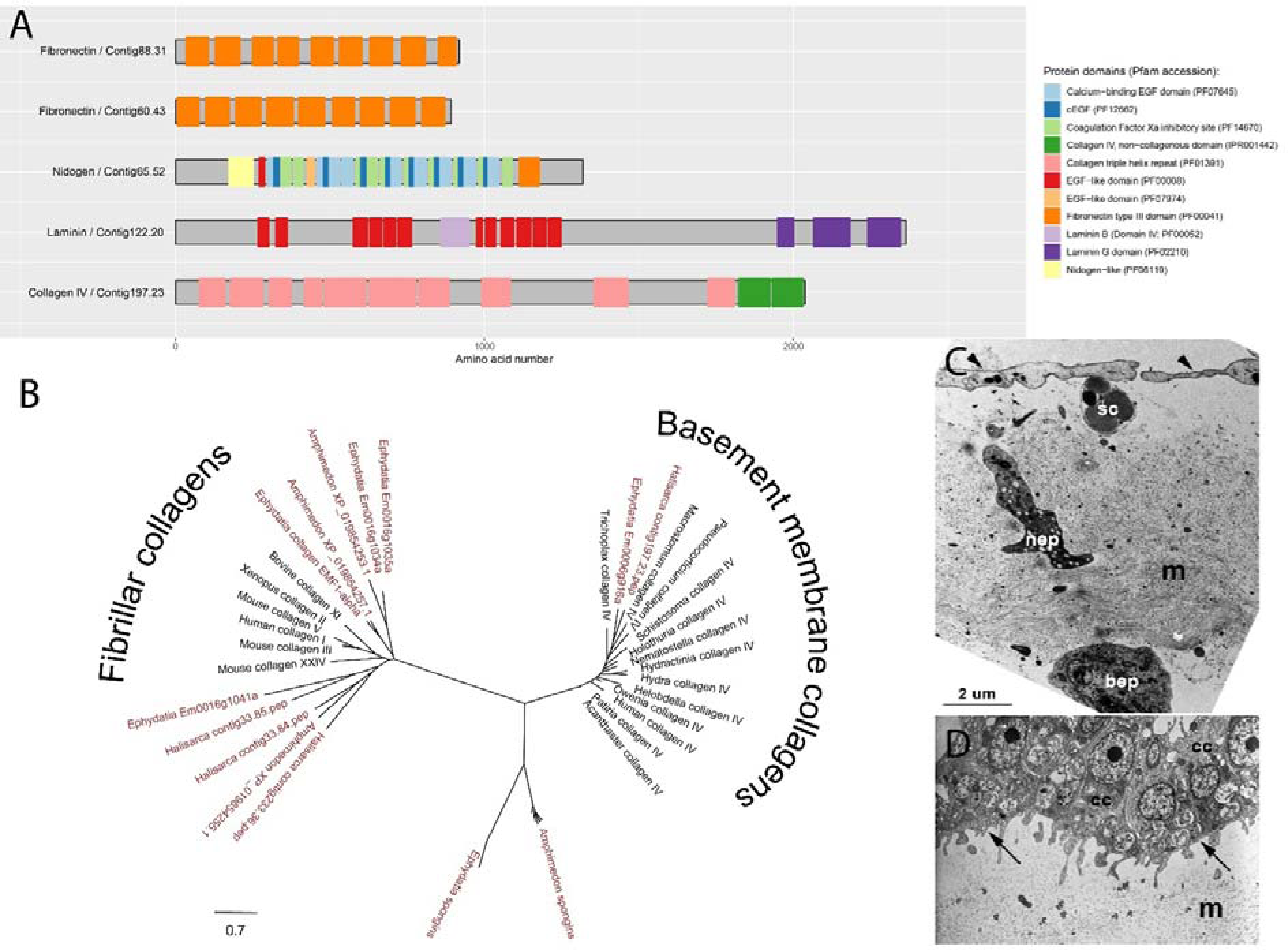
Basement membrane components in *H. dujardinii* genome. (A) Domain architecture components of key proteins evaluated by HMMER against Pfam database. (B) Unrooted maximum likelihood tree constructed from the alignment of the C-terminal NC1 domains of type IV collagens and different fibrillar collagens from phylogenetically distant animals as well as spongins. Sponge’s sequences are in red. (C) Transmission electron microscopy of exopinacocytes, the outer layer of *H. dujardinii* cells. They are T-shaped: the cell consists of an outer part (cap, shown by the *arrowhead*), a neck (*nep*) and a cell body with a nucleus (*bep*). (D) The choanocytes (*cc*) form another epithelium-like tissue in *H. dujardinii*. Their basal surfaces (indicated by *arrows*) face the mesohyl (*m*). Fibers are visible in the mesohyl, but no structures resembling the basement membrane are observed. Scale bar is the same for both TEM images. Sponge tissues were fixed in 2.5% glutaraldehyde on 0,1M cacodylate buffer, then post-fixed in 1% OsO_4_ and embedded in EMbed-812 resin (Electron Microscopy Science) after dehydration. Sections with a thickness of 70 nm were stained with lead citrate and uranylacetate.

Type IV collagen attracted our attention in particular. The most part of the molecule of type IV collagen is occupied by a helical domain consisting of triplets of Gly-Xaa-Yaa repeats, where X and Y are often proline and hydroxyproline. At the C-terminal end of the peptide is the globular domain NC1 (non-collagenous), so named because it differs from the typical fibrillar collagen helix built of repeats. In higher Metazoa, collagens are represented by a group of proteins (humans have 22 of its orthologs), some of which form fibrils (types I-III, V, and XI) and others form nonfibrillar structures (types IV, VI, VII, VIII, IX, *etc*.; Exposito et al., 2002). When the mesohyl of sponges was studied by biochemical methods based on selective extraction of fibrillar proteins, the presence of two collagen proteins named spongins was shown ^42^. A study of the ultrastructure of sponges showed that their mesohyl contains cross-striated fibers 20-25 nm in diameter, resembling collagen fibers, and in a number of structures (cement between the spicules, inorganic skeleton of bath sponges, envelope of gemmules) spongin fibers 10 nm thick are present ^43^. The study of spongins has shown that they are short-chain collagens containing a small helical domain and an NC1 domain (). Their orthologs are common in invertebrate animals from sponges to ascidians. Phylogenetic analysis suggests that spongins are precursors of type IV collagens ^44^. Recent work divides sponge collagens into 4 types: spongins, collagen IV-like proteins, fibrillar collagens of Demospongia, and collagens of glass sponges ^2^.

Type IV collagen was previously discovered among sponges by BLAST and phylogenetic analysis in *Corticium candelabrum* (Homoscleromorpha) and *Sycon coactum* (Calcarea) (Leys and Riesgo, 2012). Experimental confirmation of the localization of this protein in the basal membrane was obtained in homoscreromorphs *Pseudocorticium jarrei* and *O. lobularis* ^3,5^.

We searched the proteome of *H. dujardinii* for homologs of collagen IV, fibrillar collagens, and spongins. As Aouacheria et al. point out, “from the collagen nomenclature, noncollagenous domains have been named purely on the basis of their position from the C-terminus of the collagen chain, that is, the most C-terminal noncollagenous regions have been defined as NC1 domains although their sequences are often unrelated” ^44^. Indeed, a number of fibrillar collagens as well as collagen IV have an NC1 domain at the C-terminal end. At the same time, the Pfam database contains two different hidden Markov models for NC1 domains, PF0140 (NC1 domain of fibrillar collagens) and PF01413 (NC1 domain of collagen IV). Using HMMER, we searched for proteins containing these domains in the proteomes of *H. dujardinii*, *A. queenslandica*, and *E. muelleri*. We were unable to identify any NC1 domain in the previously described spongins ^44^, so we used them for BLAST against the same proteomes.

A search with HMMER showed the presence of one protein with the NC1 domain of collagen IV in *H. dujardinii* and *E. muelleri*. No such proteins were detected in *A. queenslandica*. Collagens with NC1 domain PF0140, presumably fibrillar, were identified in all three species: nine in *A. queenslandica* and *E. muelleri*, and sixteen in *H. dujardinii*. Three paralogs of each were taken into phylogenetic analysis. Using BLAST, several sequences of spongin short-chain collagens were obtained from *E. muelleri* and *A. queenslandica*; these had a helical domain and a C-terminal domain that did not correspond to either PF0140 or PF01413. In *H. dujardinii*, among the top-10 BBHs to *E. muelleri* spongins ^44^, there were either collagens with a PF01410 domain or no NC1 domain at all. Sequences of collagens from different Metazoa were also taken for analysis -a complete list of collagens is given in Suppl. Table 5. To find out to which group certain sponge proteins belong, we performed phylogenetic analysis (Fig. 5B). The *H. dujardinii* and *E. muelleri* proteins with domain PF01413 belong to the clade with type IV collagens. Fibrillar metazoan collagens form a group with spongin proteins containing the PF01410 domain. Spongin short-chain collagens of *E. muelleri* and *A. queenslandica* are clustered separately from these two groups.

Leys and Riesgo considered various cases of detection in sponges of proteins similar in domain organization to one or another component of the basement membrane and the absence of their complete set in the then available genome of *A. queenslandica* ^45^. It is likely that collagen IV and laminin were in the genome of the last common ancestor of multicellular animals, and the absence of one or the other genes in sponges and comb jellies is the result of secondary loss ^46^. Here we described for the first time a complete set of basement membrane proteins in the genome of *H. dujardinii*. Despite the presence of the corresponding proteins, we were unable to find morphological features of any structures resembling the basement membrane of epithelia in the layers of pinacocytes or choanocytes (Fig. 5C, D). The presence of basement membrane proteins and enzymes ensuring the assembly of the collagen network in *H. dujardinii* suggests that either (1) the protein components of the basement membrane arose independently before its formation as components of the ECM and then, as a result of convergent evolution, formed this structure, or (2) they form a homologous structure not detectable at the electron microscopic level. This question remains to be resolved in the future using immunocytochemistry and gene expression data.

### Aggregation factor

Another large group of extracellular matrix proteins that are unique to demosponges are called aggregation factors (AF). These large extracellular glycoproteins, responsible for species-specific and allorecognition, got their name for their participation in the recognition that occurs during cell aggregation ^30,47^. At the beginning of the 20^th^ century, the phenomenon of sponge development from a suspension of cells obtained by mechanical or chemical (with the help of chelators of divalent cations) dissociation was discovered ^48^. In experiments on mixing sponge cells of two different species, the cells aggregated only with cells of their own species but not with those of another. Cells interact in this process using receptors on the surface and AFs connecting receptors of neighboring cells. Allorecognition is enabled by high levels of polymorphism and RNA editing ^49^.

AFs were well characterized biochemically in three species of Demospongiae ^30,50,51^. Five clustered AF genes were then characterized in *A. queenslandica* using BLAST ^49^. A search in the transcriptomes of 24 sponge species revealed that AFs are present only within the class Demospongiae, suggesting a monophyletic origin of these molecules. A conserved region is present at the C-terminal end of the peptide, identified by Grice et al. using multiple sequence alignment and named Wreath domain ^49^. We used the authors’ hidden Markov model of the Wreath domain and the AF identification algorithm to search for AFs in the *H. dujardinii* genome based on the presence of the Wreath and Calx-beta domains. We detected 49 transcripts encoding peptides with Wreath domain transcribed from 17 genes. Of these, 14 genes are assembled in a cluster located on contig 20, and another 3 genes are scattered in other contigs (Fig. 6A). Typical (group 1 according to Grice et al., 2017) AFs of *H. dujardinii* are assembled in a cluster. They are characterized by the presence of a signal peptide, the Wreath domain at the C-terminus, and several Calx-beta domains (Fig. 6B). Calx-beta domains bind calcium cations and are responsible for adhesion ^52,53^; their number varies from 1 to 66 in typical AFs. The number of Wreath domains varies in typical AFs from 1 to 3, and due to alternative splicing, their number may vary in isoforms of transcripts of the same gene. Some peptides also contain EGF-like, von Willebrand factor type D, and other domains. Proteins encoded by three genes not belonging to the cluster are classified as group 3 according to Grice et al. One of them contains 5 Calx-beta domains but does not contain Wreath; the second one contains Wreath domain but does not contain Calx-beta, and the third one consists almost entirely of Wreath domain.

**Fig. 6.**
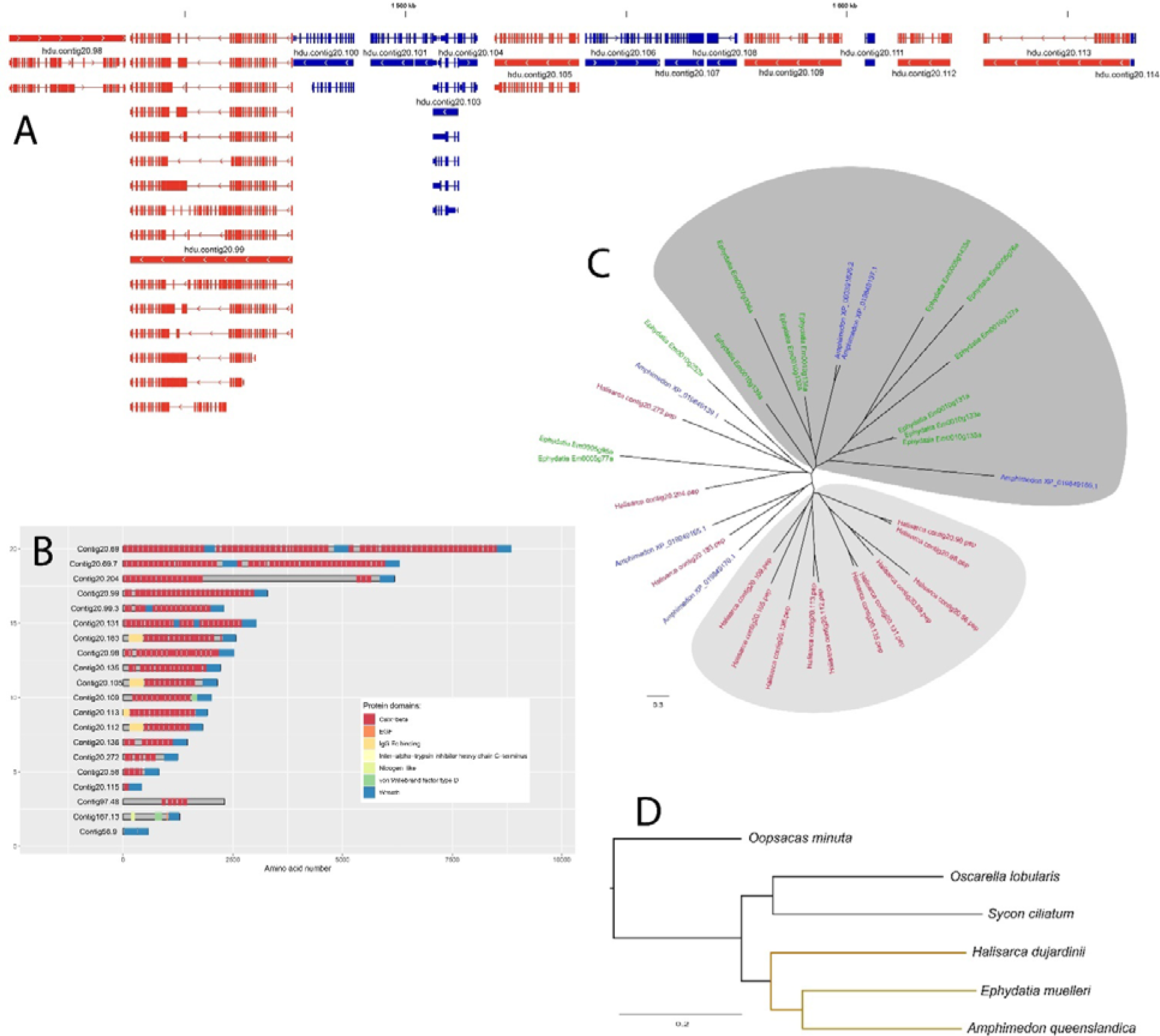
Aggregation factors of *H. dujardinii*. (A) Genome browser view of 250-kb fragment of contig 20 contains six AFs genes in *H. dujardinii*. AF genes are in red, other genes are blue. Thick blocks represent coding exons, thin blocks non-coding exons and lines introns. Small arrows on introns denote the direction of transcription. Isoforms are shown for transcripts with alternative splicing. Scale bar and position at the contig shown on top. (B) Domain composition of the AF and AF-like proteins. (C) Unrooted maximum likelihood tree inferred from the alignment of full-length AFs of *H. dujardinii*. The sequences of *E. muelleri* are shown in green, *A. queenslandica* in blue, and *H. dujardinii* in red. The clade expanded in the genome of *H. dujardinii* is highlighted in light gray; the clade combining *E. muelleri* and *A. queenslandica* sequences is highlighted in dark gray. (D) Phylogenetic tree of sponge species constructed from OrthoFinder proteome analysis (SpeciesTree_rooted). Branches of the class Demospongiae are highlighted in yellow.

We retrieved 6 proteins with the Wreath domain from the *A. queenslandica* genome and 13 ones from the *E. muelleri* genome. By aligning their full-length sequences with 14 AFs of *H. dujardinii*, an unrooted maximum likelihood tree was constructed. The alignment includes the Wreath domain and some Calx-beta domains because their number and localization are different. It is not possible to use outgroup in the tree building: a search for proteins containing the Wreath domain in Uniprot did not identify any sequences. Although the overall phylogeny of AFs cannot be unambiguously resolved using these three species, several features are notable. (1) A group of AFs shows taxon-specific expansion for *H. dujardinii* (light gray area, Fig. 6C). (2) Some genes are clearly the result of duplications, as indicated by their position in the tree and close proximity on the chromosome. (3) A large clade (dark gray area) includes proteins from *A. queenslandica* and *E. muelleri*, which within the class Demospongiae are closer to each other than to *H. dujardinii* (Fig. 6D). (4) At least one group of orthologs found in all three genomes descended from a common ancestor (*Halisarca* contig20.272, *Amphimedon* XP_019849139.1 and *Ephydatia* Em0010g252a). These data demonstrate the complexity of the evolutionary history of aggregation factors within the class Demospongiae, and their expansion in *H. dujardinii*.

The study of non-bilaterian phylogenetic lineages is of particular importance in order to understand the origin and early evolution of structures, processes, and functions in Metazoa. Phylum Porifera, located at the base of the phylogenetic tree, is a convenient ancient taxon to search for answers to such questions. Carefully assembled and annotated genomes of non-model species are still extremely scarce, although next-generation sequencing has become routine. In the present work, we presented the results of analyzing the assembly of the demosponge *Halisarca dujardinii* genome made with Oxford Nanopore long reads. In the genome, we analyzed genes encoding components of basement membrane and aggregation factors, proteins responsible for calcium-dependent adhesion and recognition. From the analysis, it is clear that even within the class Demospongiae, differences between well-studied species (such as *A. queenslandica, E. muelleri*, and *H. dujardinii*) are highly significant in such features as AFs gene set and the presence of basement membrane components. This demonstrates how phylum Porifera is heterogeneous. In addition, we have obtained a resource for further study of gene expression mechanisms by modern methods (scRNA-seq, ChIP-seq) in dynamic processes such as development and regeneration.

## Acknowledgements

We thank the resource centers ‘Computer Centre SPbU’ and ‘Molecular and Cell Technologies’ of St Petersburg State University for technical support, the Educational and Research Station of SPbU “Belomorskaia”. The research was supported by the Russian Science Foundation grant No. 22-74-00042, https://rscf.ru/project/22-74-00042/.

## References

1. Schultz, D. T. et al. Ancient gene linkages support ctenophores as sister to other animals. Nature 618, 110–117 (2023).

2. Ehrlich, H., Wysokowski, M., Żółtowska-Aksamitowska, S., Petrenko, I. & Jesionowski, T. Collagens of Poriferan Origin. Marine Drugs 16, 79 (2018).

3. Boute, N. et al. Type IV collagen in sponges, the missing link in basement membrane ubiquity. Biology of the Cell 88, 37–44 (1996).

4. Gazave, E. et al. No longer Demospongiae: Homoscleromorpha formal nomination as a fourth class of Porifera. in Ancient Animals, New Challenges (eds. Maldonado, M., Turon, X., Becerro, M. & Jesús Uriz, M.) 3–10 (Springer Netherlands, Dordrecht, 2011). doi:10.1007/978-94-007-4688-6_2.

5. Vernale, A. et al. Evolution of mechanisms controlling epithelial morphogenesis across animals: new insights from dissociation-reaggregation experiments in the sponge Oscarella lobularis. BMC Ecology and Evolution 21, 160 (2021).

6. Busch, K. et al. Biodiversity, environmental drivers, and sustainability of the global deep-sea sponge microbiome. Nat Commun 13, 5160 (2022).

7. Carrier, T. J. et al. Symbiont transmission in marine sponges: reproduction, development, and metamorphosis. BMC Biology 20, 100 (2022).

8. Sipkema, D. et al. Large-scale production of pharmaceuticals by marine sponges: Sea, cell, or synthesis? Biotechnology and Bioengineering 90, 201–222 (2005).

9. Ereskovsky, A., Borisenko, I. E., Bolshakov, F. V. & Lavrov, A. I. Whole-Body Regeneration in Sponges: Diversity, Fine Mechanisms, and Future Prospects. Genes 12, 506 (2021).

10. Srivastava, M. et al. The Amphimedon queenslandica genome and the evolution of animal complexity. Nature 466, 720–726 (2010).

11. Belahbib, H. et al. New genomic data and analyses challenge the traditional vision of animal epithelium evolution. BMC Genomics 19, (2018).

12. Fortunato, S. et al. Genome-wide analysis of the sox family in the calcareous sponge Sycon ciliatum: multiple genes with unique expression patterns. EvoDevo 3, 14 (2012).

13. Francis, W. R., et al. The Genome Of The Contractile Demosponge Tethya wilhelma And The Evolution Of Metazoan Neural Signalling Pathways. BioRxiv doi: 10.1101/120998 (2017).

14. Kenny, N. J. et al. Tracing animal genomic evolution with the chromosomal-level assembly of the freshwater sponge Ephydatia muelleri. Nat Commun 11, 3676 (2020).

15. Santini, S. et al. The compact genome of the sponge Oopsacas minuta (Hexactinellida) is lacking key metazoan core genes. BMC Biology 21, 139 (2023).

16. Wörheide, G. et al. Chapter One-Deep Phylogeny and Evolution of Sponges (Phylum Porifera). in Advances in Marine Biology (eds. Becerro, M. A., Uriz, M. J., Maldonado, M. & Turon, X.) vol. 61 1–78 (Academic Press, 2012).

17. Sambrook, J., Russell, D. W., Irwin, C. A. & Janssen, K. A. Molecular Cloning: A Laboratory Manual. (Cold Spring Harbor Laboratory Press, 2001).

18. Baril, T., Galbraith, J. & Hayward, A. Earl Grey: a fully automated user-friendly transposable element annotation and analysis pipeline. 2022.06.30.498289 Preprint at 10.1101/2022.06.30.498289 (2023).

19. Haas, B. J. et al. Automated eukaryotic gene structure annotation using EVidenceModeler and the Program to Assemble Spliced Alignments. Genome Biology 9, R7 (2008).

20. Stanke, M. et al. AUGUSTUS: ab initio prediction of alternative transcripts. Nucleic Acids Res 34, W435–W439 (2006).

21. Lomsadze, A., Burns, P. D. & Borodovsky, M. Integration of mapped RNA-Seq reads into automatic training of eukaryotic gene finding algorithm. Nucleic Acids Res 42, e119 (2014).

22. Supek, F., Bošnjak, M., Škunca, N. & Šmuc, T. REVIGO Summarizes and Visualizes Long Lists of Gene Ontology Terms. PLOS ONE 6, e21800 (2011).

23. Blin, N. & Stafford, D. W. A general method for isolation of high molecular weight DNA from eukaryotes. Nucleic Acids Res 3, 2303–2308 (1976).

24. Mayer, A. M. S. et al. Marine pharmacology in 2018: Marine compounds with antibacterial, antidiabetic, antifungal, anti-inflammatory, antiprotozoal, antituberculosis and antiviral activities; affecting the immune and nervous systems, and other miscellaneous mechanisms of action. Pharmacol Res 183, 106391 (2022).

25. Adameyko, K. I. et al. Conservative and Atypical Ferritins of Sponges. International Journal of Molecular Sciences 22, 8635 (2021).

26. Erpenbeck, D. et al. First evidence of miniature transposable elements in sponges (Porifera). Hydrobiologia 687, 43–47 (2012).

27. Fernandez-Valverde, S. L., Calcino, A. D. & Degnan, B. M. Deep developmental transcriptome sequencing uncovers numerous new genes and enhances gene annotation in the sponge Amphimedon queenslandica. BMC Genomics 16, (2015).

28. Borisenko, I., Adamski, M., Ereskovsky, A. & Adamska, M. Surprisingly rich repertoire of Wnt genes in the demosponge Halisarca dujardini. BMC Evolutionary Biology 16, 123 (2016).

29. Hodor, P. G., Illies, M. R., Broadley, S. & Ettensohn, C. A. Cell-substrate interactions during sea urchin gastrulation: Migrating primary mesenchyme cells interact with and align extracellular matrix fibers that contain ECM3, a molecule with NG2-like and multiple calcium-binding domains. Developmental Biology 222, 181–194 (2000).

30. Fernàndez-Busquets, X. & Burger, M. M. Cell adhesion and histocompatibility in sponges. Microscopy Research and Technique 44, 204–218 (1999).

31. McGregor, L. et al. Fraser syndrome and mouse blebbed phenotype caused by mutations in FRAS1/Fras1 encoding a putative extracellular matrix protein. Nat Genet 34, 203–208 (2003).

32. Shafeghati, Y., Kniepert, A., Vakili, G. & Zenker, M. Fraser syndrome due to homozygosity for a splice site mutation of FREM2. Am J Med Genet A 146A, 529–531 (2008).

33. Exposito, J. Y. & Garrone, R. Characterization of a fibrillar collagen gene in sponges reveals the early evolutionary appearance of two collagen gene families. Proc Natl Acad Sci U S A 87, 6669–6673 (1990).

34. Vallet, S. D. et al. Computational and experimental characterization of the novel ECM glycoprotein SNED1 and prediction of its interactome. Biochem J 478, 1413–1434 (2021).

35. Barqué, A. et al. Knockout of the gene encoding the extracellular matrix protein SNED1 results in early neonatal lethality and craniofacial malformations. Dev Dyn 250, 274–294 (2021).

36. Argraves, W. S., Deák, F., Sparks, K. J., Kiss, I. & Goetinck, P. F. Structural features of cartilage matrix protein deduced from cDNA. Proc Natl Acad Sci U S A 84, 464–468 (1987).

37. Kiss, I. et al. Structure of the gene for cartilage matrix protein, a modular protein of the extra-cellular matrix. Exon/intron organization, unusual splice sites, and relation to alpha chains of beta 2 integrins, von Willebrand factor, complement factors B and C2, and epidermal growth factor. J Biol Chem 264, 8126–8134 (1989).

38. McShane, A. et al. Mucus. Curr Biol 31, R938–R945 (2021).

39. Pajic, P. et al. A mechanism of gene evolution generating mucin function. Science Advances 8, eabm8757 (2022).

40. Pozzi, A., Yurchenco, P. D. & Iozzo, R. V. The nature and biology of basement membranes. Matrix Biol 57–58, 1–11 (2017).

41. Exposito, J., Cluzel, C., Garrone, R. & Lethias, C. Evolution of collagens. Anat. Rec. 268, 302– 316 (2002).

42. Gross, J., Sokal, Z. & Rougvie, M. Structural and chemical studies on the connective tissue of marine sponges. J Histochem Cytochem 4, 227–246 (1956).

43. Garrone, R. Phylogenesis of Connective Tissue. Morphological Aspects and Biosynthesis of Sponge Intercellular Matrix. (John Wiley & Sons, Hoboken, NJ, USA, 1978).

44. Aouacheria, A. et al. Insights into Early Extracellular Matrix Evolution: Spongin Short Chain Collagen-Related Proteins Are Homologous to Basement Membrane Type IV Collagens and Form a Novel Family Widely Distributed in Invertebrates. Molecular Biology and Evolution 23, 2288–2302 (2006).

45. Leys, S. P. & Riesgo, A. Epithelia, an Evolutionary Novelty of Metazoans. J. Exp. Zool. (Mol. Dev. Evol.) 318, 438–447 (2012).

46. Renard, E., Le Bivic, A. & Borchiellini, C. Origin and Evolution of Epithelial Cell Types. In Origin and Evolution of Metazoan Cell Types (CRC Press, 2021).

47. Moscona, A. A. Cell aggregation: properties of specific cell-ligands and their role in the formation of multicellular systems. Dev Biol 18, 250–277 (1968).

48. Wilson, H. V. On some phenomena of coalescence and regeneration in sponges. Journal of Experimental Zoology 5, 245–258 (1907).

49. Grice, L. F. et al. Origin and evolution of the sponge aggregation factor gene family. Molecular Biology and Evolution 34, 1083–1099 (2017).

50. Müller, W. E., Koziol, C., Müller, I. M. & Wiens, M. Towards an understanding of the molecular basis of immune responses in sponges: the marine demosponge Geodia cydonium as a model. Microsc Res Tech 44, 219–236 (1999).

51. Wiens, M. et al. Innate immune defense of the sponge Suberites domuncula against bacteria involves a MyD88-dependent signaling pathway. Induction of a perforin-like molecule. J Biol Chem 280, 27949–27959 (2005).

52. Fernàndez-Busquets, X., Körnig, A., Bucior, I., Burger, M. M. & Anselmetti, D. Self-recognition and Ca2+-dependent carbohydrate-carbohydrate cell adhesion provide clues to the cambrian explosion. Mol Biol Evol 26, 2551–2561 (2009).

53. Hilge, M., Aelen, J. & Vuister, G. W. Ca2+ regulation in the Na+/Ca2+ exchanger involves two markedly different Ca2+ sensors. Mol Cell 22, 15–25 (2006).

