## Supplementary Figures 1-4 for "First draft genome assembly and characterization of sponge *Halisarca dujardinii* reveals key components of basement membrane and broad repertoire of aggregation factors"

Supplementary Figure 1. Frequency spectra of k-mer (k=21) in natural (A) and logarithmic (B) scales, and the results of modeling the observed k-mer frequency distribution. The peak of the distribution with a coverage of about 200 corresponds to the haploid genome. The k-mer counting was performed using the jellyfish v2.3.0.

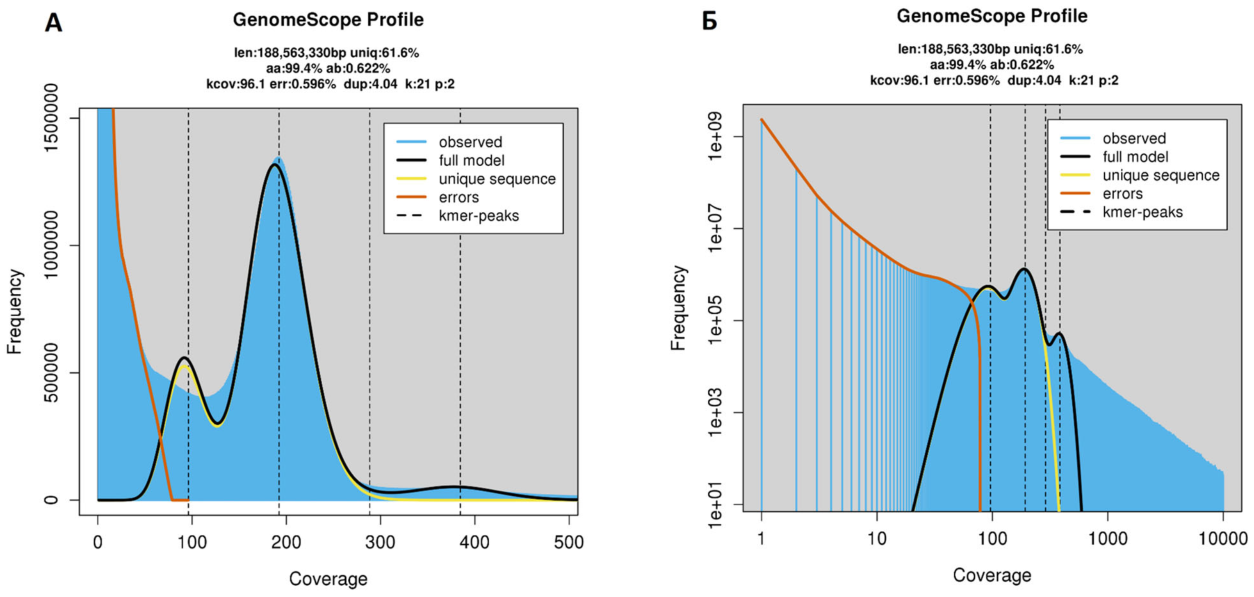


Supplementary Figure 2. Histogram of coverages of individual contigs by reads obtained by Flye assemblyer.

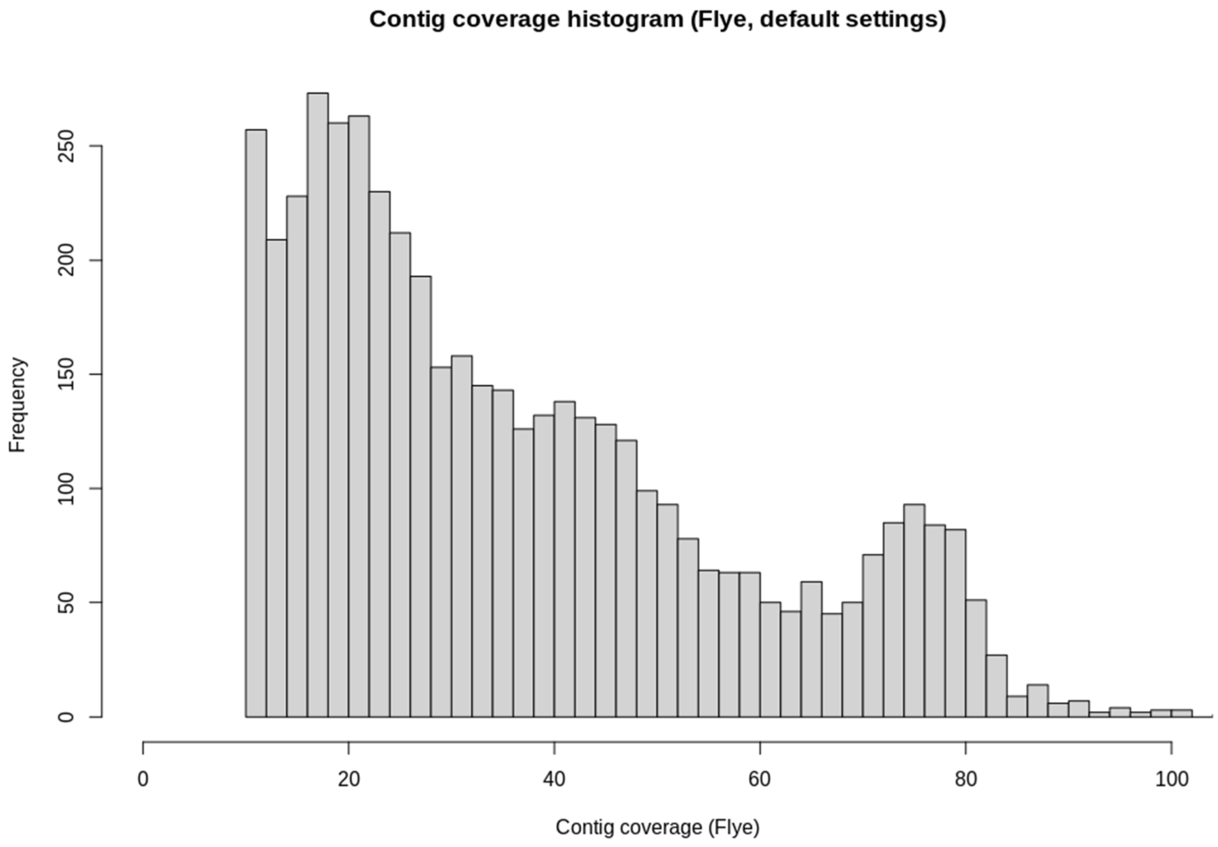

Supplementary Figure 3. Results of polishing genomic assemblies using long reads and the Racon software.

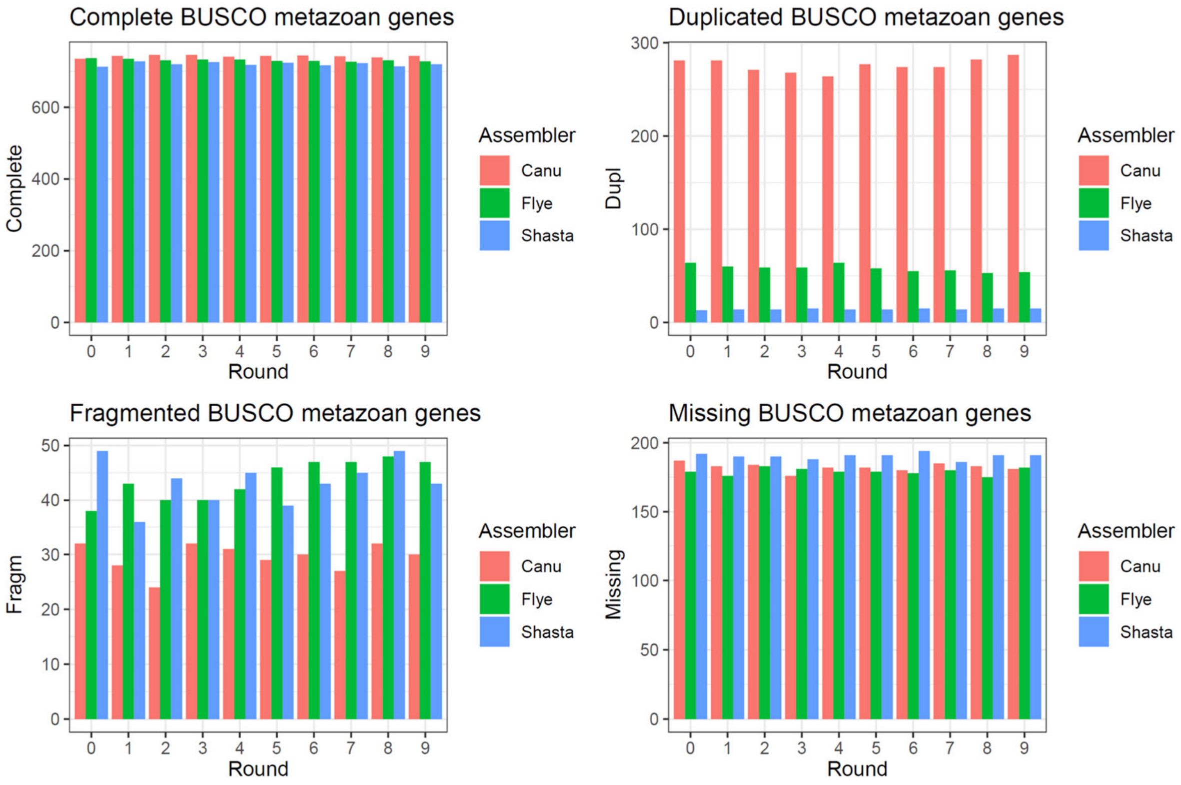

Supplementary Figure 4. (A) Orthology analysis performed on sponges, cnidarians, placozoans and Deuterostomia. (B) GO enrichment of sequences unique for *H. dujardinii*. GO terms in category “Cell compartment” are indicated inside cells, and cell square proportional to p-value.

A

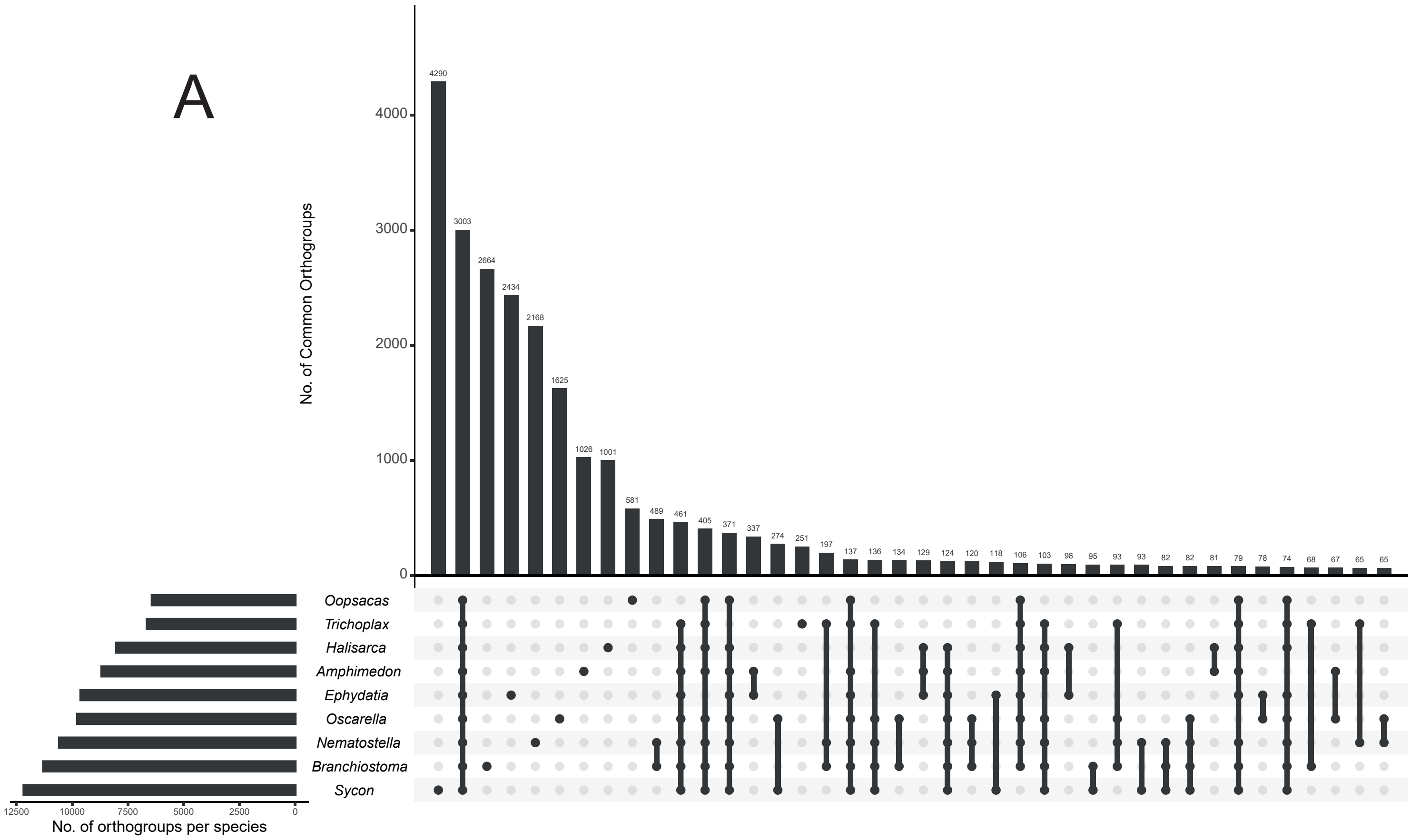


B

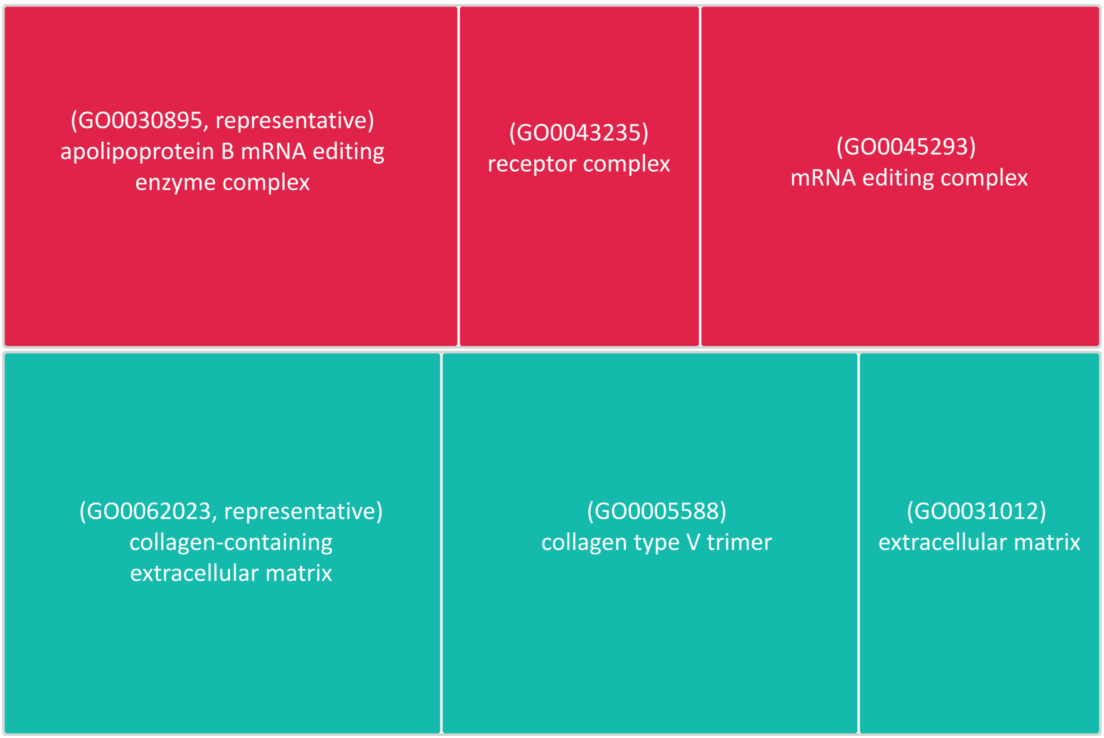
